# The regulation of mammalian maternal-to-embryonic transition by Eukaryotic translation initiation factor 4E

**DOI:** 10.1101/2020.01.16.908582

**Authors:** Yan Li, Xuefeng Huang, Jianan Tang, Xu Ji, Miao Liu, Lu Chang, Jing Liu, Yihua Gu, Changgen Shi, Wuhua Ni, Hui-juan Shi, Chris O’Neill, Xingliang Jin

## Abstract

Genetic and inhibitor studies show expression of eukaryotic translation initiation factor 4E (eIF4E) was required for the successful maternal-to-embryonic transition of mouse embryos. eIF4E was in both gametes and in the cytoplasm and pro-nuclei soon after fertilization, and at each stage of early development. Knockout (*Eif4e*^−/−^) by *PiggyBac (PB) [Act-RFP]* transposition caused peri-implantation embryonic lethality due to the failure of embryos to form a pluripotent epiblast. Maternal stores of eIF4E supported development up to the 2-4-cell stage after which new expression occurred from both alleles. Inhibition of the maternally acquired stores of eIF4E (4EGI-1 inhibitor) resulted in a developmental block at the 2-4-cell stage. 4E-BP1 is a hypophosphorylation-dependent negative regulator of eIF4E. mTOR activity was required for 4E-BP1 phosphorylation and inhibiting 4EGI-1 retarded embryo development. eIF4E expression and activity is regulated at key embryonic transitions in the mammalian embryo and is essential for successful transition to embryonic control of development.

**Significance Statement:** eIF4E is recognized as the rate-limiting factor for CAP-dependent translation. This work used a combination of a gene knockout model, selective pharmacological inhibition and expression analyses to investigate the expression and function of Eif4e in the early mouse embryo. It provides compelling evidence for the essential role of Eif4E in the normal processes of early mammalian embryo development, including the formation of the pluripotent epiblast and the maternal-embryonic transition. The unexpected evidence for a growth deficit in mice hypomorphic for Eif4e will be a key area of future investigation. It also provides for the first time a powerful demonstration of the utility of the *PB [Act-RFP]* transposon mouse model for analyzing the molecular regulation of early mammalian embryo development.

## Introduction

Reproduction in all metazoan requires the conversion of the terminally differentiated gametes into the totipotent cells of the early embryo. The earliest stages of embryo development are under the control of transcripts and protein inherited from the gametes, primarily the oocyte. The transition from the maternal to embryonic control of embryo development is accompanied by the generation of a new transcriptome and proteome. In mammals, the oocyte and zygote are transcriptionally inert until embryo genome activation (EGA) (1, 2). In the mouse, new transcription is initiated in zygotes and definitive embryonic transcription commences in the 2-cell embryo (3). This is accompanied by the degradation of almost all the maternally inherited transcripts (4). Some understanding of the details of reprogramming of the proteome in the early embryo in mammals is also now emerging (5–7). Many proteins present within the oocyte are also detected in the zygote, however, a proportion are rapidly lost after fertilization (5). This is accompanied by the increased protein expression of many components of the ubiquitin/proteosome protein degradation. A range of other proteins are present at higher levels in the zygote than the oocyte and include proteins associated the citrate cycle pathway, glucan metabolism, lipid binding proteins, and fatty acid metabolism (5–7). This early round of new translation also includes components of the transcriptional machinery (8) and translational activity is required for the successful activation of the embryonic genome (9). After EGA the new transcriptome is subject to a further round of translation, however, this does not result in a strong correlation between the RNA species generated at EGA and the cellular proteome until the morula and blastocyst stages (5, 6).

The factors regulating translation in the mammalian embryo have not received detailed analysis. In eukaryotes, more than 95% of proteins are synthesized through 5’ methylguanosine (m^7^G) cap-dependent mRNA translation (10, 11). Eukaryotic translation initiation factor 4E (eIF4E) is generally considered the rate-limiting factor in translation from mRNA (12). This protein recognizes and binds to the m^7^G cap moiety on mRNA within the cytoplasm (13). eIF4E participates in the formation of a multiprotein complex that also contains eIF4A and eIF4G (10) in the formation of this complex is a rate-limiting step in cap-dependent translation (14). eIF4E binding to mRNA cap structures is inhibited by a small family of eIF4E-binding proteins (4E-BPs) in their hypo-phosphorylated state (15). However, the phosphorylation of 4E-BP dissociates it from eIF4F, allowing the formation of the eIF4E complex and initiation of cap-dependent translation (11, 16). Among these proteins, 4E-BP1 is the most abundant and is phosphorylated at multiple sites by a phosphoinositide (PI) 3-kinase/ Protein kinase B (Akt)/ mechanistic target of rapamycin kinase (mTOR)- dependent mechanism (11). eIF4E function can also be regulated by phosphorylation at Serine 209 by a p38 mitogen associated protein kinase (mitogen-activated protein kinase 14, Mapk14, also known as p38 MAPK) and MAPK signal-integrated kinase (MNK) signaling pathway (17, 18). The role of eIF4E phosphorylation (p-eIF4E) is not clear since hypo-phosphorylated eIF4E can bind the mRNA caps and stimulate translation *in vitro* (10, 19), although p-eIF-4E is reported to enhance the translation of a subset of proteins (20).

In sea urchins, fertilization triggers dissociation of eIF4E from 4E-BP allowing rapid recruitment of eIF4E into a high molecular mass complex. 4E-BPs are rapidly phosphorylated (21) and degraded following fertilization, and this involves a rapamycin-sensitive mTOR signaling pathway (22, 23). In mammalian oocytes, temporal and spatial control of translation is regulated via an mTOR–eIF4F pathway (24).

Genetic studies have shown that *Eif4e*^+/−^ mice are viable (12) yet details of the phenotype of *Eif4e-*null embryos have not to our knowledge been described in detail. In this study, we used mouse *PB* transposon induced transgenesis to produce a functional *Eif4e* knock-out. It showed that *Eif4e*^−/−^ was embryonic lethal, with lethality occurring around the time of implantation. eIF4E showed temporal and sub-cellular regulation across early embryo development and was required for successful embryonic development.

## Results

### eIF4E and p-eIF4E in the mouse preimplantation embryo and gametes

Western blot analysis showed a single band of eIF4E (~33 kDa) and phosphorylated eIF4E (p-eIF4E) (~28 kDa) in mouse oocytes and preimplantation embryos (Fig 1A). The larger size of band for eIF4E may indicated a post-translational modification, such as sumulation (25, 26). The detection level of both antigens was relatively stable relative to ACTIN levels across each developmental stage, except for lower p-eIF4E at the 2-cell stage (Fig 1A). Immunofluorescence detected abundant eIF4E and p-eIF4E in the mid-piece of sperm (Fig 1B). eIF4E was present in oocytes and all stages of preimplantation embryos (Fig 1C). The eIF4E antigen staining was evenly distributed in across the oocyte but in the in the embryo it was characterized by an accumulation within the nucleolus precursor bodies (NPB) present in both parental pronuclei in the zygotes and in 2-cell stage embryos. This was not evident from the 8-cell embryo. P-eIF4E was detected at all stages except in the 2-cell stage embryo (Fig 1D). It was evenly distributed across the cells at each of these stages of development (Fig 1 A, B, D). Unlike the native protein, p-eIF4E did not show an obvious accumulation within NPBs. This relatively pervasive presence of eIF4E and p-eIF4E in the gametes and across early stages of embryo development is indicative of a likely role in these reproductive processes while its differential localization patterns at key embryonic transitions may be indicative of active regulation.

**Figure 1.**
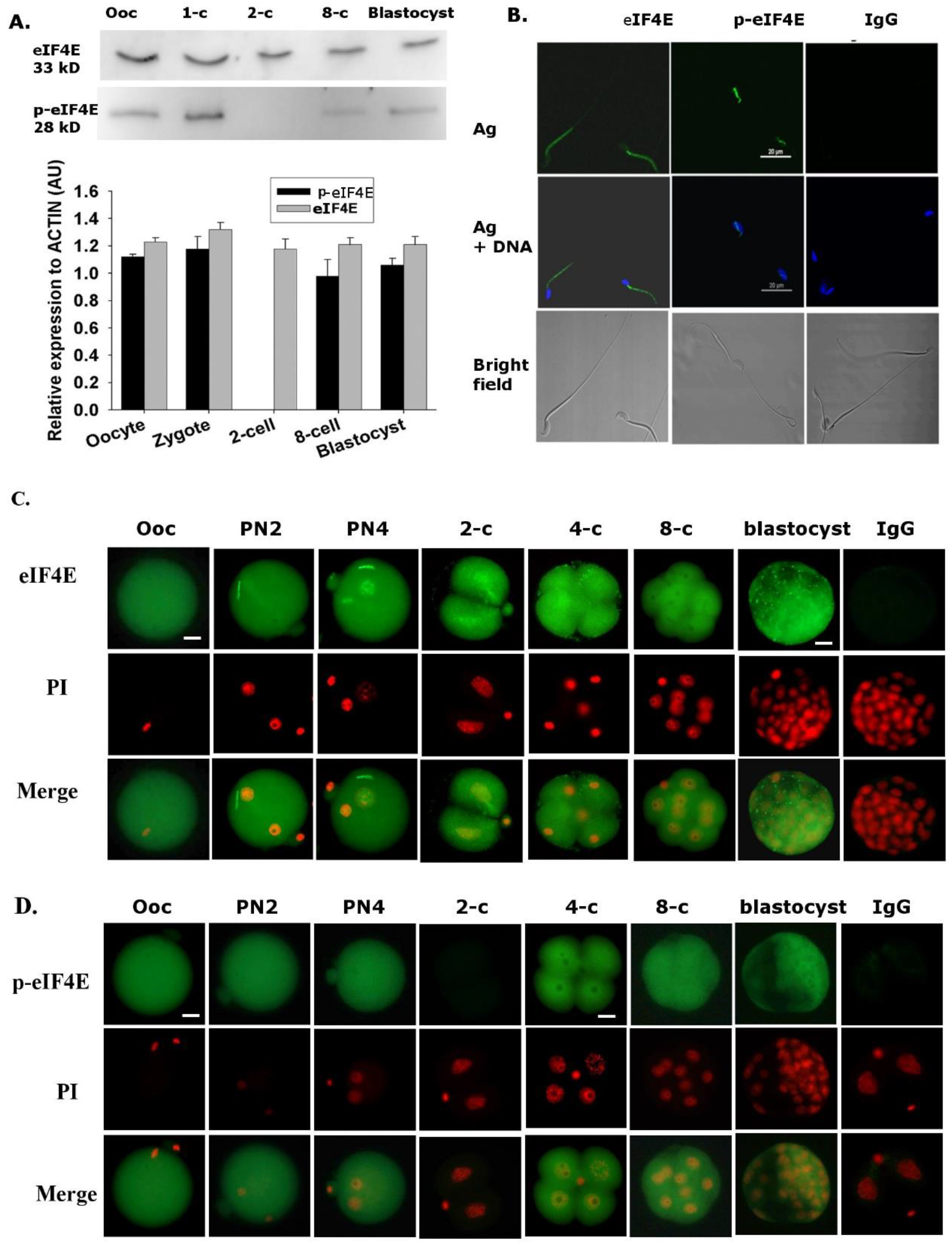
eIF4E and p-eIF4E in mouse gametes and preimplantation embryos. (A) The presented images were representative of three replicates of Western blot analysis of eIF4E and p-eIF4E antigens, and their relative fold changes to ACTIN in oocytes and preimplantation embryos. Each sample contained 50 oocytes or embryos. The molecular weights of eIF4E and p-eIF4E were 32.98 ± 0.13 kDa and 28.05 ± 0.16, respectively. (B) Confocal-fluorescent images of eIF4E and p-eIF4E with Hoechst33342 staining in mouse sperm. Negative controls staining conditions with non-reactive IgG and generated no signal in sperm. (C) (D) The presented images were whole session epi-fluorescent staining of eIF4E and p-eIF4E encountered with propidium iodine staining. The data were representative of over 30 embryos of each developmental stage. Negative controls staining conditions with non-reactive IgG and generated no signal in any of the developmental stages. Scale bar = 20 μm for images in B and = 10 μm in C and D.

### The developmental viability of Eif4e deficient mice

Cross-breeding *Eif4e*^+/−^ mice produced only *Eif4e*^+/+^ or *Eif4e*^+/−^ offspring, no *Eif4e*^−/−^ progeny were born (Table 1). *Eif4e*^+/−^ pups were fluorescent red (Fig 2 A) due to the expression of the RFP-reporter present within the *PB [Act-RFP]* construct (Fig 2B). The total number and the number per litter of *Eif4e*^+/−^ pups was smaller than expected theoretical Mendelian ratios (MR) (P < 0.01) (Table 1). At 4-weeks of age, the body weight of *Eif4e*^+/−^ pups was lower than in their *Eif4e*^+/+^ littermates (P < 0.01) (Table 1) for both sexes, but there was no significant difference between sexes within the same genotype. A number of sites with incomplete or inviable embryonic tissue were present at embryonic day 7.5 (E7.5) and for each the dissected embryonic tissue was from *Eif4e*^−/−^ embryos. At E10.5, only *Eif4e*^+/−^ and *Eif4e*^+/+^ embryos were present (Fig 2C). These results show that *Eif4e*-null embryos are not capable of development beyond early gestation while *Eif4e* hypomorphic mice were viable but had reduced postnatal growth rates.

**Table 1.**
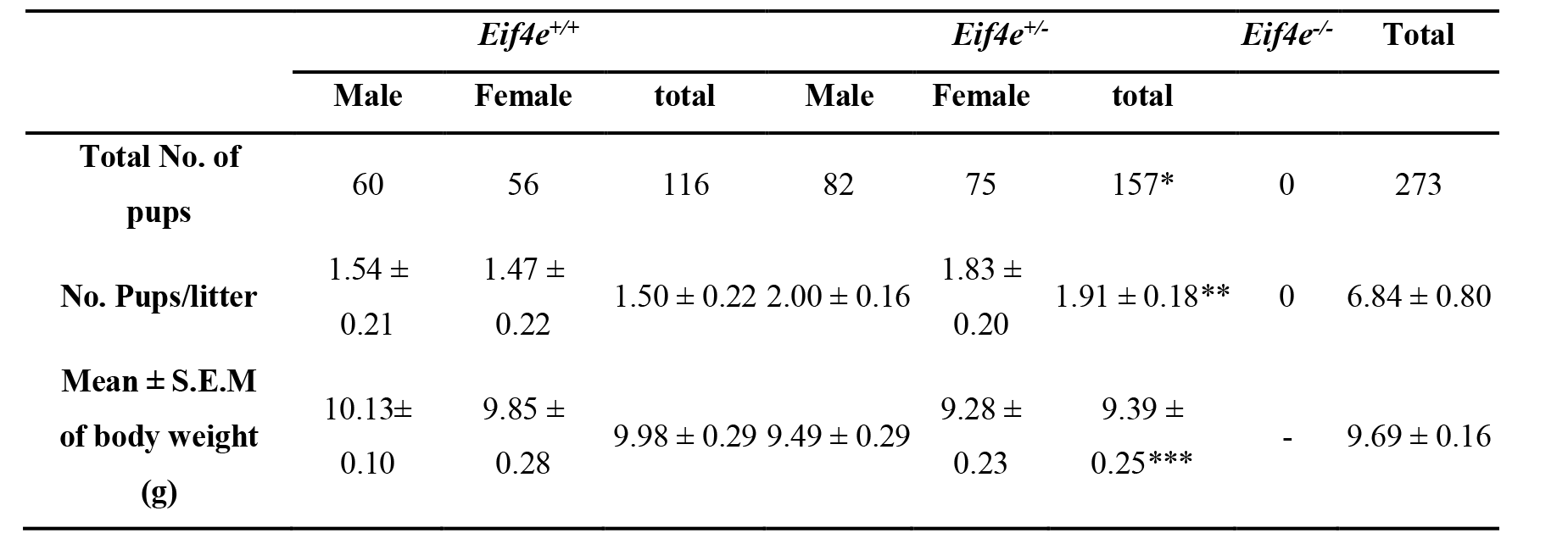
Birth outcomes of cross-mating of *PB* transposon-based *Eif4e*^+/−^ mice. Total number of pups, pup number per litter and body weight were analyzed after cross mating of *Eif4e*^+/−^ mice resulted from 39 births. Chi-square test was performed to compare the total number of pups (* p<0.01) and pup number per litter (**p<0.01) of *Eif4e*^+/−^ mice with *Eif4e*^+/+^in according to mendelian ratio (MR). Mean ± S.E.M of body weights of 4-week-old *Eif4e*^+/−^ compared with *Eif4e*^+/+^ (*** p<0.05), and between the both sexes within the same genotype (P>0.05).

|  | *Eif4e*<sup>+/+</sup> |  |  | *Eif4e*<sup>+/-</sup> |  |  | *Eif4e*<sup>-/-</sup> | Total |
| --- | --- | --- | --- | --- | --- | --- | --- | --- |
|  | Male | Female | total | Male | Female | total |  |  |
| **Total No. of pups** | 60 | 56 | 116 | 82 | 75 | 157* | 0 | 273 |
| **No. Pups/litter** | 1.54 ± 0.21 | 1.47 ± 0.22 | 1.50 ± 0.22 | 2.00 ± 0.16 | 1.83 ± 0.20 | 1.91 ± 0.18** | 0 | 6.84 ± 0.80 |
| **Mean ± S.E.M of body weight (g)** | 10.13 ± 0.10 | 9.85 ± 0.28 | 9.98 ± 0.29 | 9.49 ± 0.29 | 9.28 ± 0.23 | 9.39 ± 0.25*** | - | 9.69 ± 0.16 |

**Figure 2.**
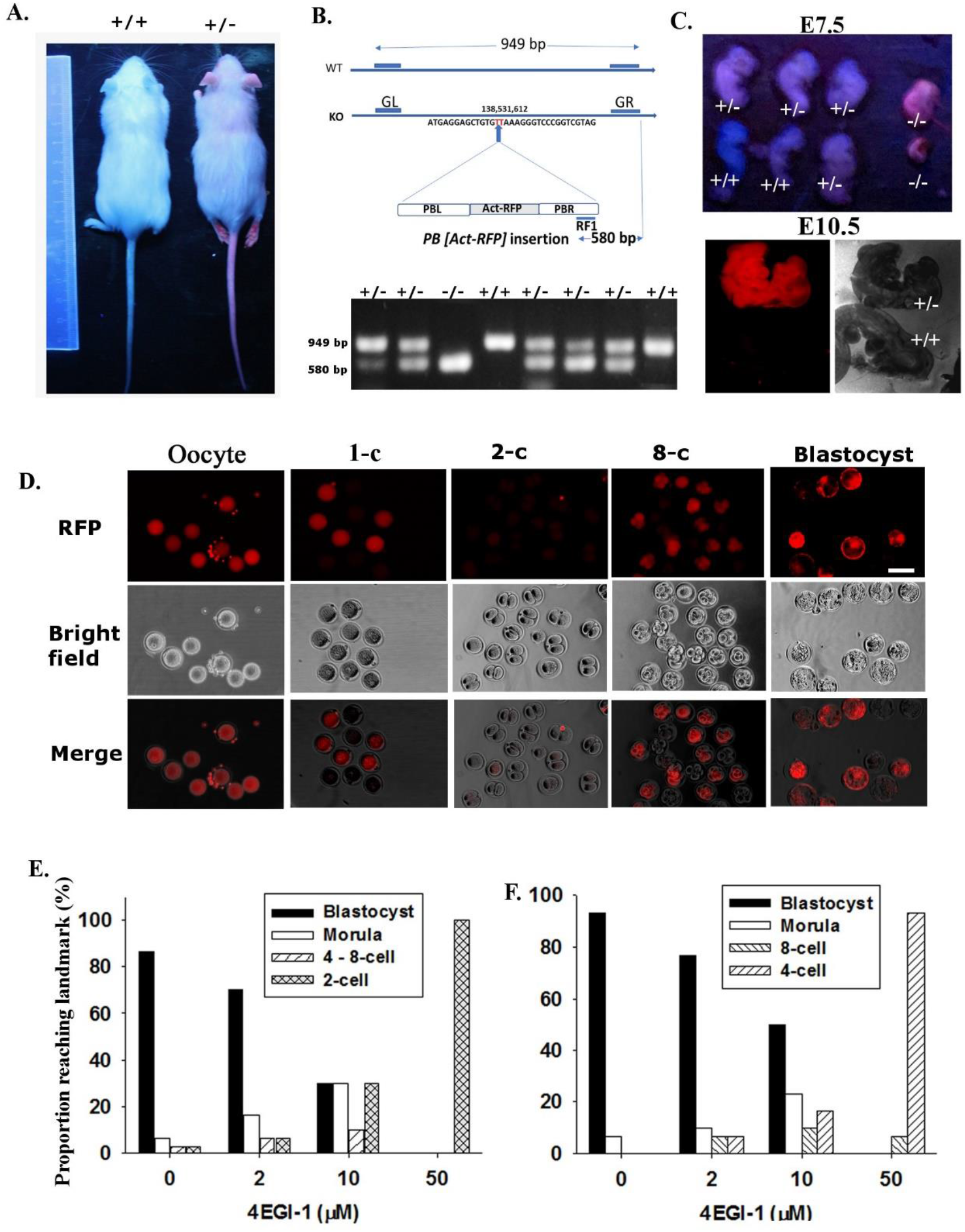
Functional knockout of *Eif4e* gene with PB transposon and pharmacological inhibition on mouse early embryo development. (A)The presented images showed the RFP^−^ *Eif4e*^+/+^ and RFP^+^ *Eif4e*^+/−^pups from *Eif4e*^+/−^ cross-mating. (B) Brief construct and gel electrophoresis analysis of *PB [Act-RFP]* transposon and its insertion position in *Eif4e* gene. Mouse *Eif4e* gene (NC_000069.6) is localized at 138,526,191 - 138,559,696 on Chromosome 3 (ENSMUSG00000028156). The insertion of *PB [Act-RFP]* transposon was constructed (48) and was posited at TTAA target site 138,531,612 of 2^nd^ intron of *Eif4*e gene. GL and GR primes detected 949 bp product representing wildtype Eif4e. The PB primer (RF1, CCTCGATATACAGACCGATAAAACACATGC) to detect this insertion was localized on the PBR which gave the 580 bp products. (C) Dissection of *Eif4e*^+/−^ females to collect embryos of E7.5 embryos and E10.5 embryos and show their RFP expression and genotyping. (D) The presented images were representative of RFP expression in oocyte and preimplantation developmental stages. (E) (F)The effects of 4EGI-1 on preimplantation development in vitro. The data were representative of three independent replicates cultured hybrid zygotes (E) or 2-cell embryos (F) in media dosed with 4EGI-1 for 96 h or 72 h, respectively. Statistical analysis was as described in the main text.

To gain an insight into the likely causes of this loss of viability of *Eif4e*-null embryos we next examined the expression of the RFP-reporter gene and early development rates of embryos derived from *Eif4e*^+/−^ x *Eif4e*^+/−^ cross-mating. All oocytes collected from the reproductive tracts of the *Eif4e*^+/−^ females were RFP-positive (RFP^+^), indicating its pre-meiotic II expression (Fig 2D). After fertilization, approximately half of the zygotes were RFP^+^ while no morphological 2-cell stage embryos collected from the reproductive tract were RFP ^+^ (Fig 2D, Table 2, suppl table1). In the 4-8-cell and blastocyst stage ~69% of embryos were RFP^+^ (Fig 2D, Table 2, suppl Table1).

**Table 2.**
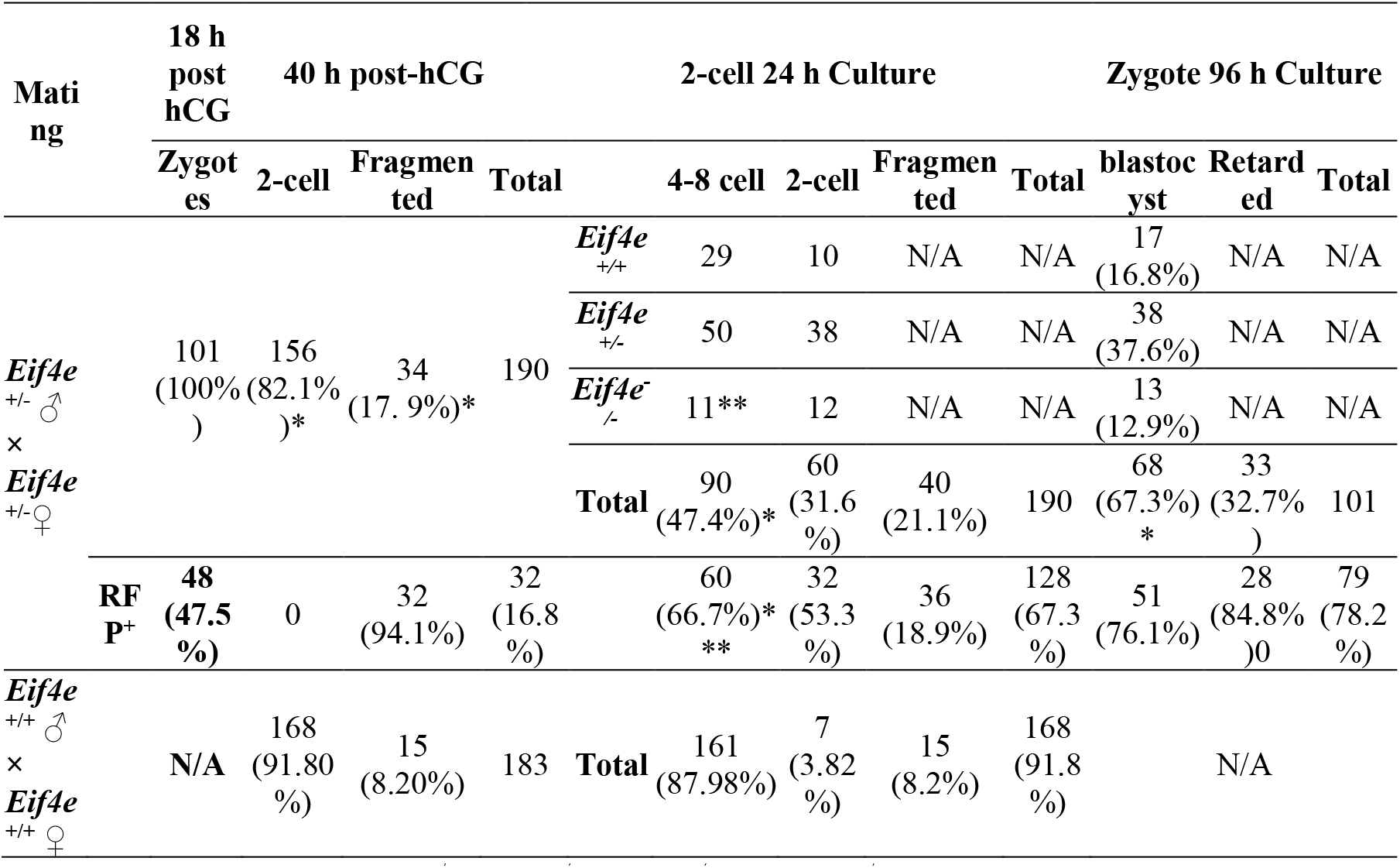
Embryo development from *Eif4e*^+/−^ ♂x *Eif4e*^+/−^ ♀ mating. The proportion of embryos *Eif4e*^+/−^ × *Eif4e*^+/−^ or *Eif4e*^+/+^♂ × *Eif4e*^+/+^♀ matings. Embryos were collected at 18h post hCG then cultured for 96h, or collected at 40h post-hCG and then cultured for a further 24h. *p < 0.01, comparing corresponding developmental stages of *Eif4e*^+/+^♂ × *Eif4e*^+/+^♀. Chi square test analyzed the rates of 4-8cell embryos (**p<0.05) and RFP^+^ (***p<0.02), according to Mendelian ratio. N/A - not available.

Approximately 18% of embryos were retarded or significantly fragmented when collected 40 h post hCG and of these 94% were RFP^+^ (Table 2) (it was not possible to reliably genotype these cells). This rate of fragmentation was significantly higher than observed in embryos collected from wildtype crosses (Table 2). Culturing the 2-cell embryos for a further 24 h resulted in approximately 50% developing to the 4-8-cell stage and of these 66.7% were RFP^+^. Individual genotyping of these 4-8-cell stage embryos showed that 67.7% were either heterozygous or homozygotes, thus by the 4-8-cell stage approximately 98% of embryos carrying ACT-RFP had regained the capacity for the expression of RFP. By contrast only 53% of embryos that were retarded after 24h culture (remaining at the 2-cell stage) were RFP^+^ while genotyping showed 83% of the retarded embryos were either heterozygous or homozygotes (p > 0.05) (Table 2). Genotyping of individual blastocysts cultured for 96 h from the zygote stage did not show any skew from expected Mendelian distribution (p >0.05). All heterozygous or homozygotes embryos were RFP^+^ while no wild-type embryos expressed RFP^+^. The results showed that eIF4E is a maternal product that is lost from the embryo progressively after fertilization. It then becomes progressively re-expressed from the late 2-cell stage, the stage at which definitive transcription from the embryonic genome is known to be initiated (27). This analysis also shows that the rapid degradation of maternal stores and the re-activation of RFP expression allows RFP detection to serve as a reliable marker of embryos past the 8-cell stage that carry the mutant allele.

Analysis of the distribution of embryos between each of the expected genotypes showed that at the 4-8-cell stage fewer *Eif4e*^−/−^ were detected than expected (p < 0.02) (Table 2) but at the blastocyst stage this skew was not significant (p > 0.05). A number of embryos were retarded in development (having not reached the expected developmental landmark of 4-8cell at 48 h post-hCG or normal morphological blastocysts after 96h culture) and of these 94% (P < 0.05 compared with expected 75% and 85% (P > 0.05) were RFP^+^ respectively (Table 2). The results show that while there may be some loss of *Eif4e*^−/−^ embryos prior to the blastocysts stage, most embryos lacking the gene could form morphological blastocysts.

The significant stores of eIF4E observed in the gametes and carried over into the zygote and the high rate of embryonic loss by the 2-cell stage point to a possible role for the protein prior to the onset of definitive embryonic genome activation. The presence of the gametic stores do not allow this question to be explored with this genetic model so the effect of selective pharmacological inhibition of eIF4E (4EGI-1; K_D_ = 25 μM (16)) on this early stage of embryo development was assessed. This antagonist blocks eIF4E binding to eIF4G preventing the formation of active complexes (16). Embryos collected at either the zygote or 2-cell stage were cultured in the media supplemented with a dose range of 4EGI-1. All doses caused significant developmental block (p < 0.001). At the highest dose the drug caused a complete developmental block of zygotes (blocked at the 2-cell stage) and of 2-cells (blocked at the 4-cells stage) (Fig 2 E, F). By contrast the presence of 50 μM 4EGI-1 did not affect fertilization by either IVF or ICSI (Table 3) but continued culture in the presence of 4EGI-1 did prevent development of the resulting zygotes to the morphological 2-cell embryo. When fertilized eggs derived in 4EGI-1 medium were transferred into control medium the blastocyst formation rate was rescued compared to controls (Table 3). This result indicates eIF4E activity from gametic stores is not required for each gamete’s capacity for fertilization but was necessary for the normal processes involved in the maternal to embryonic transition.

**Table 3.**
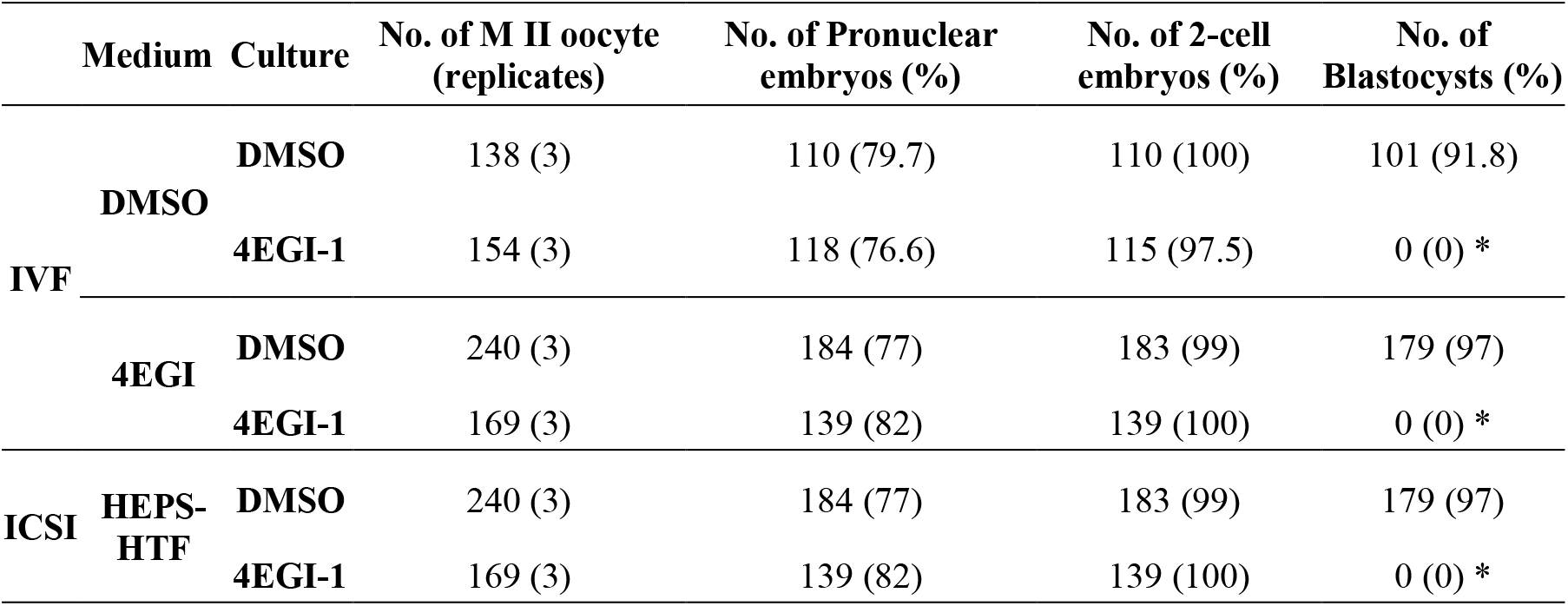
The effect of eIF4E inhibitor on IVF and development. IVF was performed in the medium containing 0.05% DMSO or 50 μM 4EGI-1. The produced embryos with pronuclei were further cultured in medium containing 0.05% DMSO or 50 μM 4EGI-1. ICSI was performed in HEPS-HTF and surviving oocytes were cultured in media with 0.05% DMSO or 50 μM 4EGI-1. The number and percentage of pronuclear embryos, 2-cell embryos, and blastocysts were recorded during development. The data were representative of three independent replicates, and each treatment included at least 40 oocytes. P*<0.001, compared the rates of blastocyst in DMSO groups.

Given that *Eif4e*^−/−^ embryos had some capacity to develop to morphological blastocysts yet no viable null embryos were detected on E7.5 we assessed the capacity of the *Eif4e*^−/−^ blastocysts to form normal outgrowths possessing a pluripotent epiblast in vitro. This showed that *Eif4e*^−/−^ blastocysts commonly had reduced expansion and either failed to produce outgrowths or produced very poor outgrowths which were smaller in size (Fig 3 A, B) and generally failed to produce an Oct3/4-positive pluripotent epiblast compared to heterozygous or wildtype embryos (Fig 3C, D, Table 4). The results point to a requirement for the new eIF4E stores produced embryo genome activation for the formation of a pluripotent epiblast in the implanting embryo.

**Table 4.**
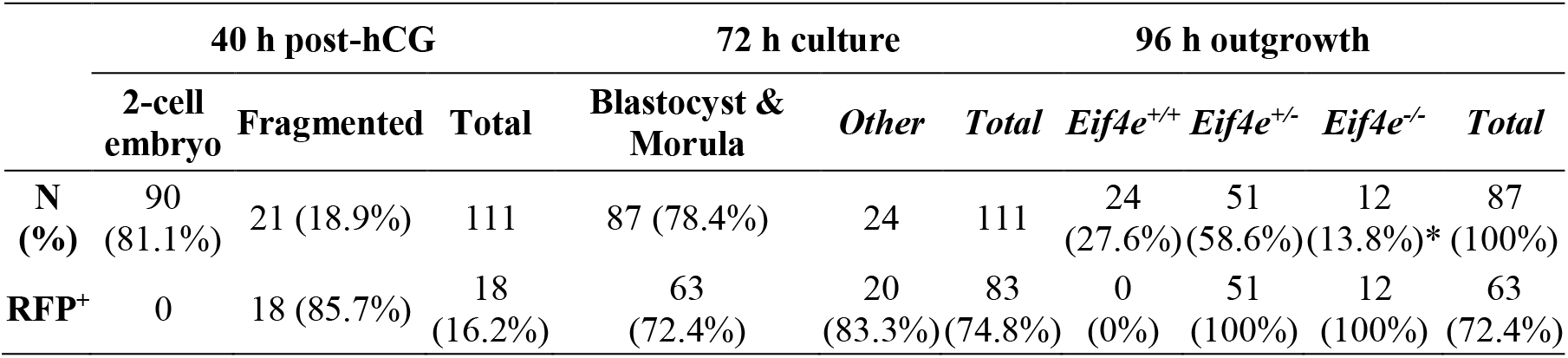
Developmental viability and RFP expression of 2-cell embryos from *Eif4e*^+/−^ ♂ x *Eif4e*^+/−^ ♀ mating. The rate of RFP+ embryos and developmental ability of 2-cell embryos collected from *Eif4e*^+/−^ cross mating. The derived morula and blastocysts were cultured for 96 h in ESC medium and genotyping was done in individual embryos. * p< 0.01 using Chi square test comparing blastocyst rates according to Mendelian ratio.

**Figure 3.**
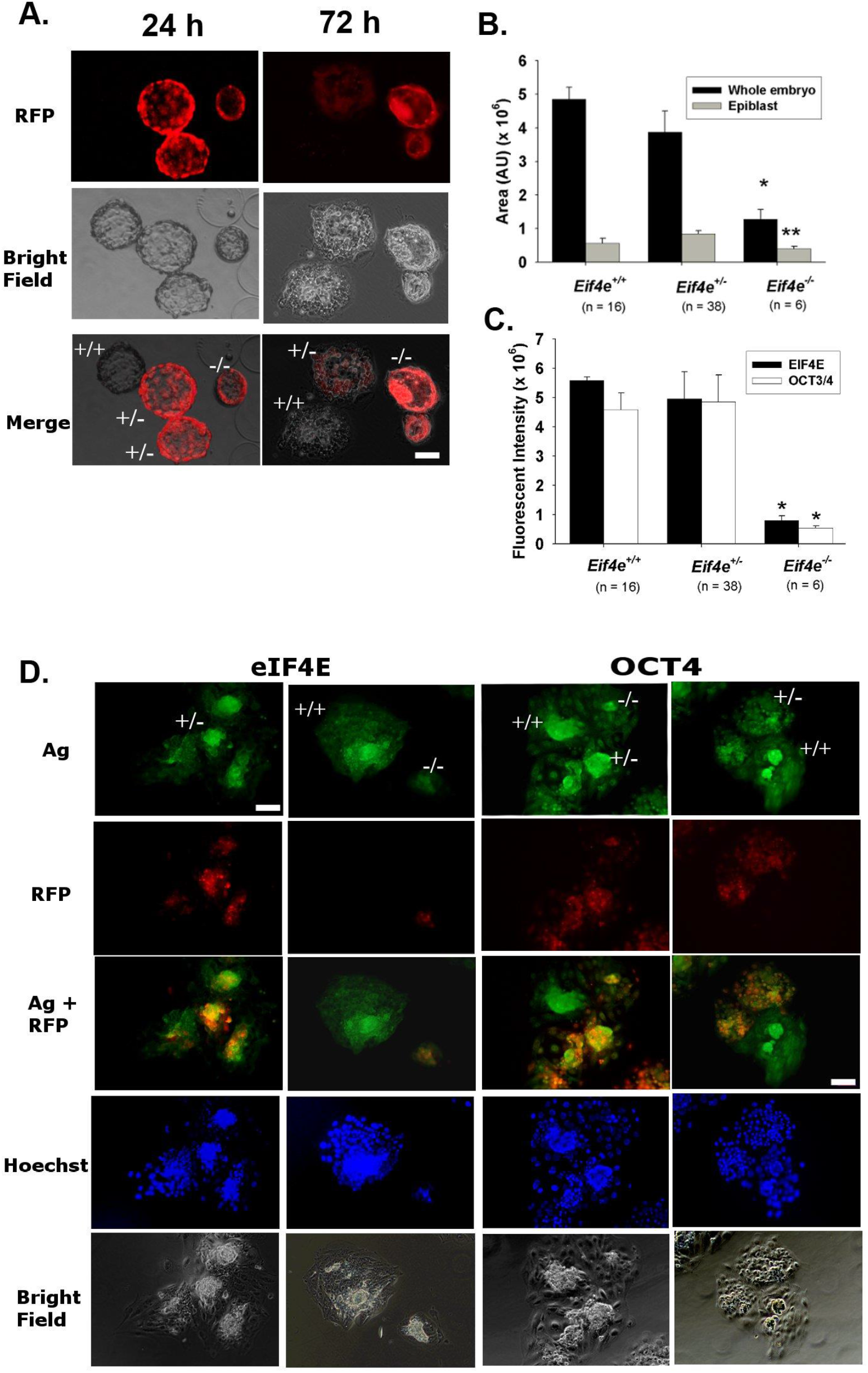
Outgrowth viability of blastocysts from *Eif4e*^+/−^ cross mating. Morphologically blastocysts derived from the culture of 2-cell embryos from *Eif4e*^+/−^ cross mating were cultured for 96 h in the media that were verified to support further growth of blastocyst. (A) The presented images of RFP expression in outgrowth embryos labelled with their individual genotype. (B) The area of whole embryo and epiblast and (C) the fluorescent intensity of eIF4E and OCT4 staining of the embryos after 96 h culture are displayed (* P < 0.001 or ** P < 0.01). The results were representative of 58 total embryos from five mates (Table 4). (D) The presented images were whole session epi-fluorescent staining of OCT4 and eIF4E, RFP reporter and nuclei (Hoechst 33342). (A, D). Scale bar = 20 μm for all images. Ag = antigen, i.e. eIF4E or OCT4.

### Gametic contributions of *Eif4e*

Immunolocalization analysis showed that eIF4E was readily detected within the oocyte and sperm and inhibitor studies indicate a possible role for these stores in early embryo development. We next utilized the RFP-reporter construct to assess the relative roles of each of these gametic stores to the expression of eIF4E across preimplantation embryo development. To achieve this, we mated either *Eif4e*^+/−^ males with *Eif4e*^+/+^ females or *Eif4e*^+/−^ females with *Eif4e*^+/+^ males. Both these reciprocal crosses produced the expected mendelian ratio of *Eif4e*^+/−^ pups (p>0.05), although there were fewer wildtype males than expected in both crosses (p < 0.05). There was no effect of pedigree on the number of pups per litter. In both crosses, the body weight of *Eif4e*^+/−^ pups was slightly lower than *Eif4e*^+/+^ pups (p<0.05) and this was not different between sexes within the same genotype (Tables 5).

**Table 5.**
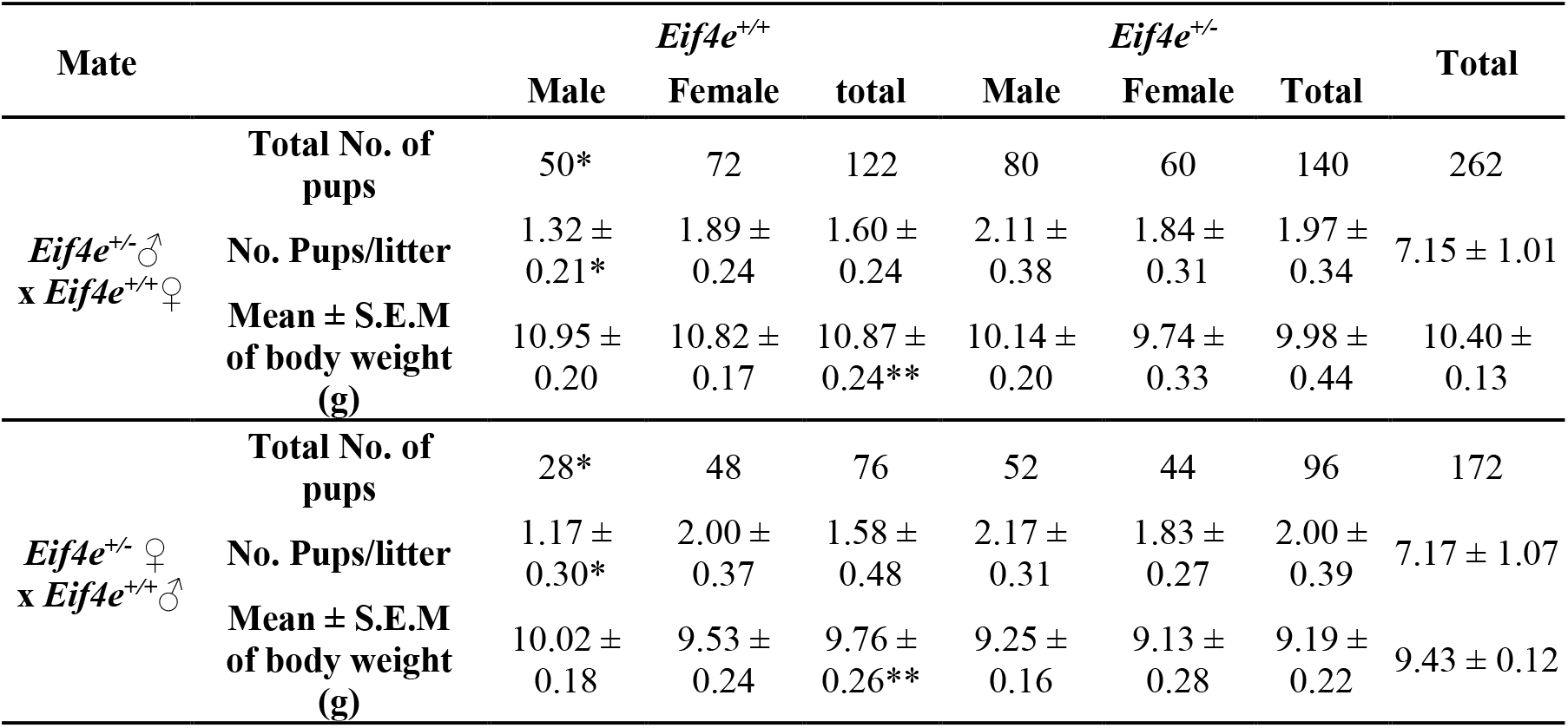
Birth outcomes of reciprocal mating of *Eif4e*^+/+^ X *Eif4e*^+/−^. The birth outcomes of 38 *Eif4e*^+/−^ ♀x *Eif4e*^+/+^ ♂matings and 24 *Eif4e*^+/−^♂ x *Eif4e*^+/+^♀ matings. Chi-square test was performed to compare the total number of pups and pup number per litter of *Eif4e*^+/+^ mice with *Eif4e*^+/−^ (*p>0.05), and *Eif4e*^+/+^ males with *Eif4e*^+/+^ females (*p <0.05). Mean ± S.E.M of body weights of was compared between both genotypes (**p<0.05), and between both sexes within the same genotype (p>0.05).

All zygotes resulting from *Eif4e*^+/−^ males with *Eif4e*^+/+^ crosses failed to express RFP. These zygotes were cultured and the onset of RFP expression monitored across early development. After 48h culture approximately a third of the resulting 4-8 cell stage embryos were RFP^+^ and by 96 h approximate 50% blastocyst were RFP^+^ (Table 6, Fig 4-A). By contrast, approximately half of zygotes produced by *Eif4e*^+/+^♂ × *Eif4e*^+/−^ ♀ crosses were RFP^+^ and from the late 2-cell embryo to the blastocyst stage the proportion stabilized around 50% RFP^+^ (Table 6, Fig 4 B). These results show maternal stores of eIF4E primarily account for the eIF4E detected within the early stages of development and that transcription from both parental alleles commences soon after the onset of definitive embryonic genome activation. The results do not provide evidence for a meaningful contribution of the eIF4E present in the sperm.

**Table 6.**
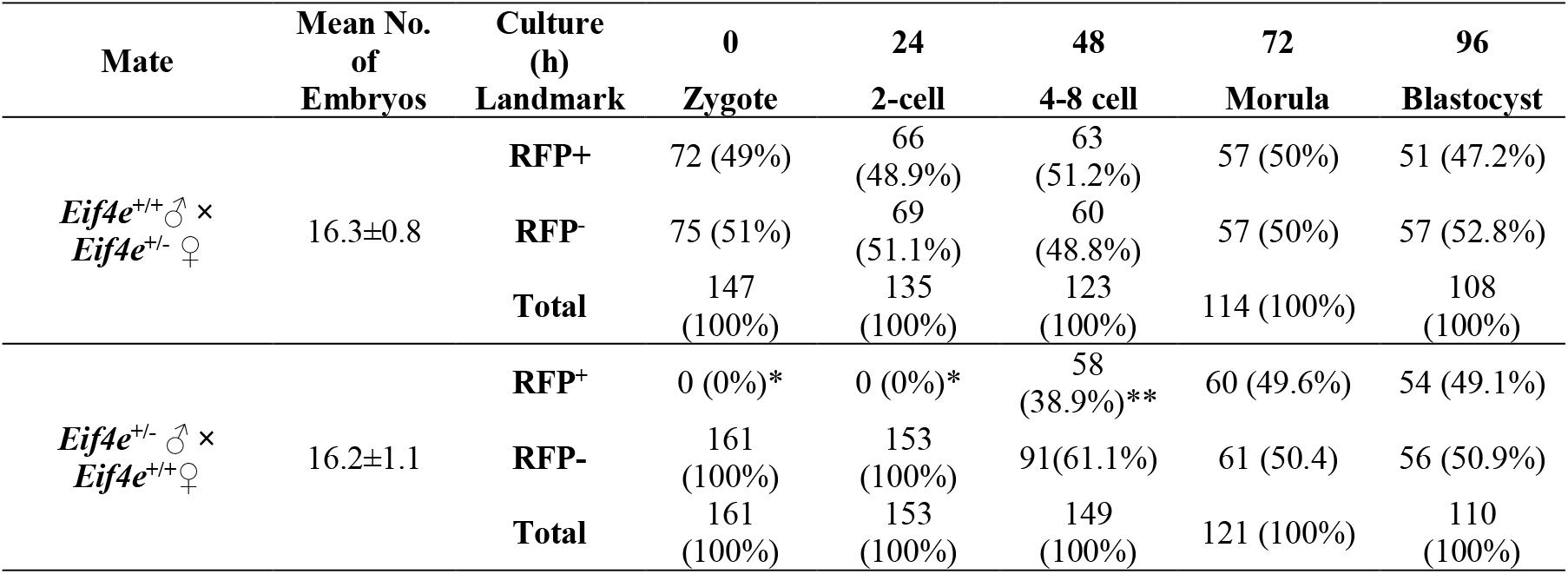
The proportion of RFP expression in developmental landmarks of the culture of the zygote from reciprocal bred mating. The rate of RFP^+^ embryos and developmental ability was monitored during in vitro development of zygotes from 9 pairs of *Eif4e*^+/+^♂ × *Eif4e*^+/−^ ♀ mates and 10 pairs of Eif4e^+/−^ ♂ × Eif4e^+/+^ ♀ mates. Chi square test analyzed the rates of RFP^+^ embryos (*p<0.001 and **p <0.01) according to Mendelian ratio.

**Figure 4.**
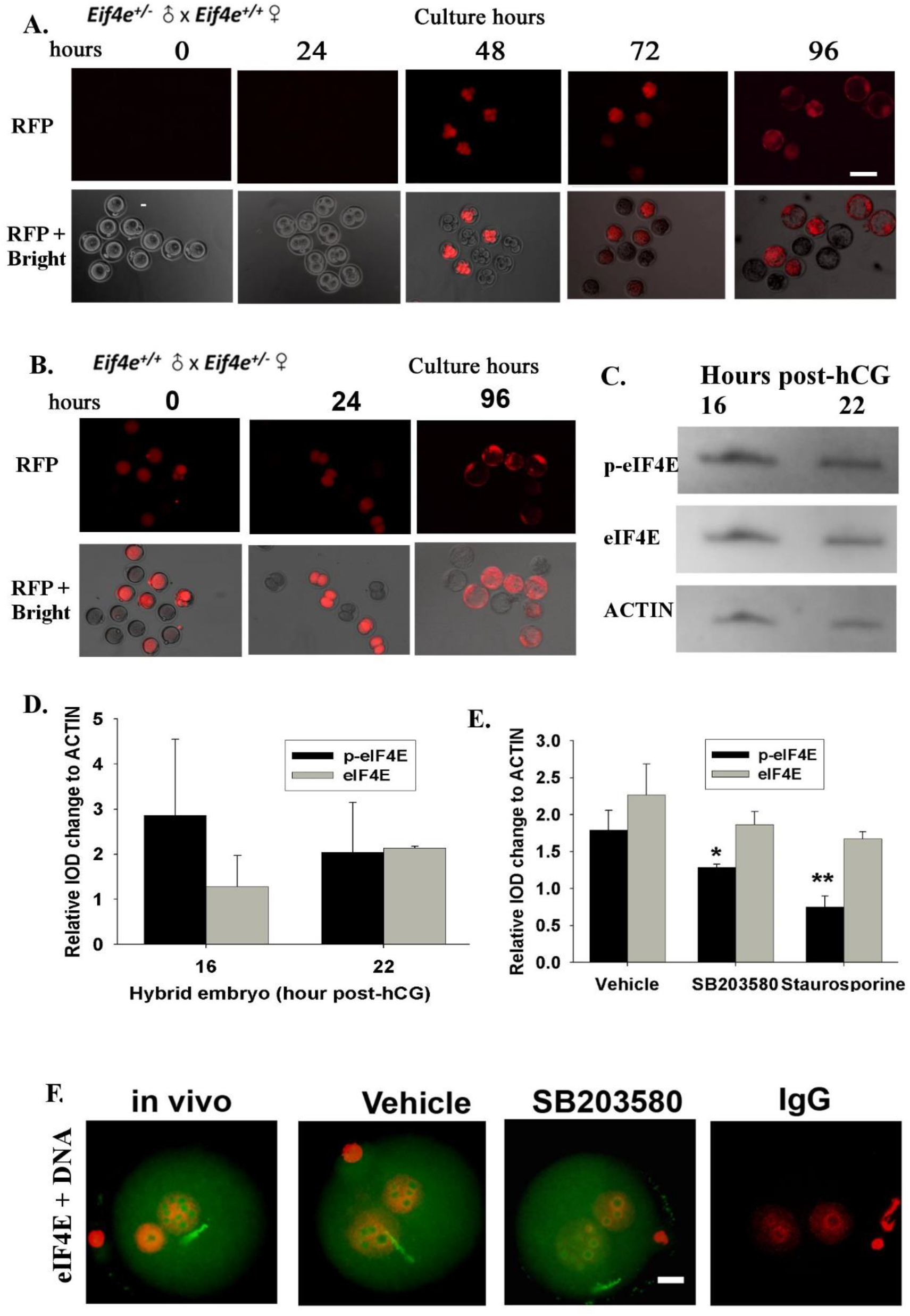
Gametic contributions of *Eif4e* and the regulation of eIF4E in zygotes. (A) (B)The expression of RFP reporter and developmental outcomes from reciprocal mating. The presented images of RFP expression in zygote cultures from *Eif4e*^+/−^ ♂ x *Eif4e*^+/+^♀ and *Eif4e*^+/+^ ♂ x *Eif4e*^+/−^ ♀ up to 96 h. Representative images of three independent replicates, where at least 20 embryos were included for each replicate. Scale bar = 50 μm. (C) (D) western blot analyzed eIF4E and p-eIF4E in hybrid zygotes, (E) the effects by SB203580 and Staurosporine, and (F) the whole-session immunolocalized images of eIF4E in SB203580 zygotes. The data were representative of three independent replicates for western blot analysis of eIF4E and p-eIF4E in hybrid zygotes in vivo and cultured zygotes treated with SB203580 or Staurosporine. * p <0.05, **p < 0.01, compared to vehicle.

### Regulation of EIF4E in the early embryo

A characteristic feature of the localization of the eIF4E antigen is its significant accumulation within the NBPs (Fig 1C). Western blot analysis showed similar levels of eIF4E and pSer209-eIF4E in the PN3 (16h post hCG) and PN5 (22 h post hCG) (Fig 4C, D). Treatment of zygotes from 16 h post hCG with either 25 nM Staurosporine (a broad spectrum protein kinase inhibitor with some preference for Protein Kinase C) or 10 μM SB203580 (a selective P38 MAPK inhibitor) (28) both reduced the levels of eIF4E phosphorylation but did not affect the total eIF4E levels (Fig 4E) in the resulting PN5 stage zygotes. Immunolocalization showed SB203580 did not affect the subcellular localization of eIF4E staining, with the accumulation within the NPBs still evident after this treatment (Fig 4F). These results show that the maintenance of maternal stores and NBP localization of eIF4E over this period of development was independent of the actions of a range of protein kinases but that maintenance of cellular levels of pSer209-eIF4E was dependent of the actions of kinases, including P38 MAP kinase.

Another striking feature of the actions of the protein was the failure to detect the Ser209 p-eIF4E antigen within the 2-cell embryo. This was surprising given the evidence that inhibition of eIF4E in the 2-cell caused a complete developmental block. While phosphorylation has some control functions, a dominant regulator of eIF4E is eukaryotic translation initiation factor 4E-binding protein 1 (4E-BP1). In its hypo-phosphorylated state 4E-BP1 acts as a translation repressor protein and is a negative regulator of eIF4E-RNA complex formation(11). Its phosphorylation by mTOR reverses this repressor activity, allowing eIF4E to undergo normal binding with 5’ m^7^G cap mRNA(16). Immunolocalization showed 4E-BP1 to be present throughout the cytoplasm and nucleoplasm of the 2-cell embryo and was a clear enrichment at the nuclear periphery and peri-nuclear regions while its level was restricted within the NBPs (Fig 5A). The phosphorylated form (pT45 4E-BP1+2+3) was also widely distributed across each of the cells of the 2-cell embryo. Its staining was characterized by a striking enrichment at the cleavage furrow (Fig 5B). We demonstrated that mTOR and Ser2448 p-mTOR (Suppl Fig1) were present through-out the 2-cell embryo and in each other stage of preimplantation embryo development. Treatment of embryos with a selective inhibitor of mTOR (PP242; IC50 = 8 nM) (29) had no effect on the level or localization of 4E-BP1 in the 2-cell but caused a large loss of pT45 4E-BPs-signal across the cells (Fig 5D). Treatment of embryos with this drug also caused a dose-dependent retardation to embryo development (Supplementary Fig1). S6K1 is another important phosphorylation target of mTOR and the inhibition of its phosphorylation (pT389-S6K1) by PP242 served as a control demonstrating the selectivity of the drug’s actions (Fig 5C, D). An unexpected finding was that the inhibition of eIF4E by 4EGI-1 also blocked phosphorylation of 4E-BP1 and S6K1, suggestive of a feedback mechanism operating between eIF4E and mTOR activity in the 2-cell. It is concluded that the activity of mTOR results in the phosphorylation of 4E-BP1 in the 2-cell embryo, a result expected to favour eIF4E activity (Fig 5), and that mTOR activity was necessary for normal embryo development.

**Figure 5.**
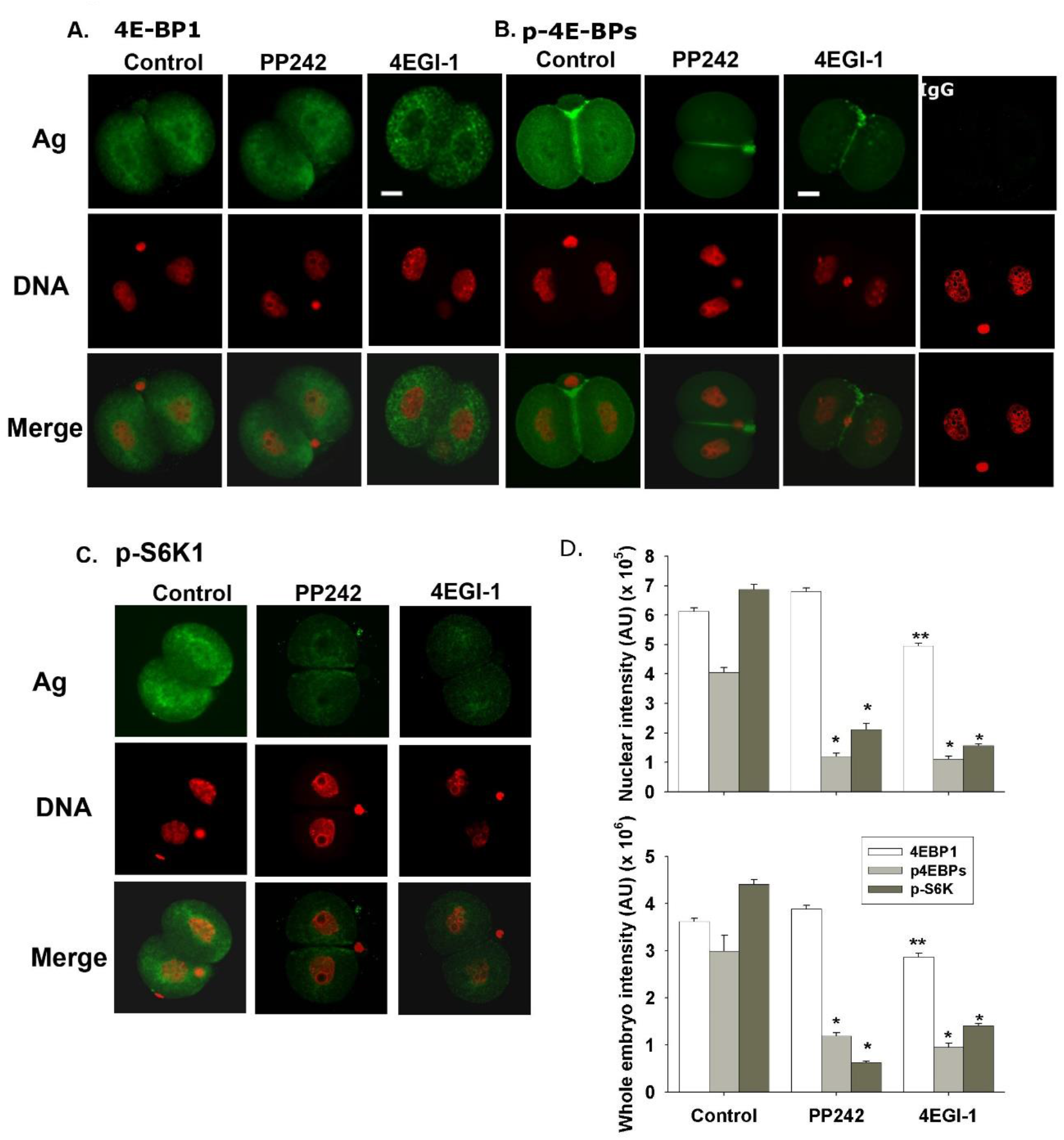
mTOR-related 4E-BP1 and S6K1 signaling in mouse 2-cell embryos. Representative confocal images of p4E-BPs (A), 4E-BP1 (B) and p-S6K1 (C) in 2-cell embryos that were derived from zygotes after treatment with 100 nM PP242 and 50 μM 4EGI-1. Scale bar = 10 μm for all images of embryo. The mean ± S.E.M of the fluorescence intensity for each antigen in the nuclei and whole embryos were displayed (D). The data were representative of three independent replicates and each treatment included at least 10 embryos. * P <0.001 and ** P < 0.01, comparing the corresponding treatments to the control.

## Discussion

PB-mediated transgenesis provides a highly efficient strategy for generating mice with loss of gene function while allowing simultaneous monitoring of a fluorescence reporter-protein expression patterns (30). This model was used to generate a loss of *Eif4e. Eif4e-*null embryos were inviable, with lethality occurring soon after embryo implantation. While *Eif4e*^−/−^ embryos resulting from *Eif4e*^+/−^ cross-mating were capable of development to morphological blastocysts they failed to reliably show blastocyst outgrowth and did not form a pluripotent epiblast. *Eif4e*^+/−^ embryos were viable but resulted in progeny with a lower post-natal body weight.

The survival of *Eif4e*^−/−^ embryos to the blastocyst stage was possibly due to the persistence of maternal protein within the early embryo. Analysis of the RFP-reporter showed that protein is carried over from the oocyte but not the spermatozoon. New expression was then initiated after definitive EGA and this occurred from both maternally and paternally inherited alleles (31). The complete developmental block of zygotes or 2-cell embryos caused by treatment with a selective eIF4E inhibitor indicates that the stores carried over from the oocyte were necessary for development. Given the expression of eIF4E from both alleles at EGA it was interesting that *Eif4e*^+/−^ embryos developed normally past the implantation stage. This result is consistent with observations that mammalian cells express levels of eIF4E that are in excess of that required for normal cell function. Furthermore, *Eif4e*^+/−^ cells showed no apparent change in global protein synthesis (12, 32). It was noteworthy that *Eif4e*^+/−^ progeny had a smaller post-natal body weight which shows that some haploinsufficiency occurred in the heterozygous state. After *Eif4e*^+/−^ X *Eif4e*^+/−^ mating there was a small deficit in the number of *Eif4e*^+/−^ progeny but this did not occur after *Eif4e*^+/−^ X *Eif4e*^+/+^ reciprocal mating, indicating that haploinsufficiency was influenced by some unexpected complexity in the interactions between the gamete carrying the null-allele and the maternal genotype.

This genetic and pharmacological evidence for the necessary actions of eIF4E at critical transitions during early embryo development are consistent with similar discoveries in non-mammalian model species, such as in sea urchins(22, 23), zebrafish(33) and *Drosophila(34)*. The pervasive detection of both eIF4E and its phosphorylated form across preimplantation stages of embryo development also supports its important roles in development. eIF4E is not normally present with nucleoli of somatic cells (35) so its accumulation within the NPBs of the zygote is interesting. NPBs are not considered to be functional nucleoli in the zygote. They lack fibrillar centres and the granular and dense fibrillar components seen in functional nucleoli.

Furthermore they seem not to undertake ribosome production (36) (37). Yet NPB function is essential for normal embryonic development indicating non-canonical roles for these structures (36). Their primary role seems to be related to the re-programming and reorganisation of centromeres and pericentric satellites into chromocentres (38). Analysis of the role of eIF4E in the reprogramming of chromatin structure and organisation in the zygote will be of interest.

Incorporation of eIF4E into the complexes required for the initiation of translation is regulated by its phosphorylation as well as through binding of inhibitory proteins (39). eIF4E phosphorylation occurs in somatic cells via a P38 Map kinase /MAP kinase (MNK) mediated pathway (40, 41),(32) and we show that P38 activity was required to maintain p-eIF4E levels in zygotes. Interestingly, while p-eIF4E was present in most stages of development tested it was not detected in 2-cell embryos and it has also been shown that 4E‐BP1 becomes dephosphorylated in the porcine embryo after fertilisation (39). The roles of eIF4E phosphorylation are not fully defined. It is not required for normal cell growth or development in some models (42) although it does seem to enhance the translation from some mRNA species (20) indicating that the dynamic changes in eIF4E phosphorylation around the time of EGA warrants further investigation.

The binding of 4E-BP1 to eIF4E can block phosphorylation of eIF4E (41, 43). The interaction of 4E-BP1 with eIF4E is in turn negatively regulated by 4E-BP1 phosphorylation via a PI3-kinase/AKT/mTOR signaling pathway (44). Autocrine trophic ligands activate the PI3K/AKT pathway soon after fertilization and this activity is essential for the normal development and survival of the early embryo (45–47). Here we show that mTOR and its phosphorylated form are also present in the 2-cell embryo. Inhibition of mTOR activity inhibited phosphorylation of 4E-BP1 in the zygote and also blocked embryo development. The results indicate that one role for autocrine trophic signaling pathways in the early embryo is to foster the activity of eIF4E.

This study shows eIF4E, an essential component of the translation initiation complex, is a critical maternal effect product that is required for the development of the early mammalian embryo. It’s presence at the critical embryonic transitions of EGA and the first rounds of cellular differentiation places it as a key regulator of the transition of maternal-to-embryonic control of development. A detailed analysis of the factors regulating translation in the embryo is an essential precondition to understanding the normal development of the embryo.

## Materials and Methods

### Animal experiments

Animal experiments were approved by and conducted according to ethics guidelines from relevant research institutes and universities. FVB mice and heterozygous *Eif4e*^+/−^ mice were obtained from the Institute of Developmental Biology and Molecular Medicine of Fudan University (Shanghai, China). *Eif4e*^+/−^ mice were generated by random germline transposition of *PiggyBac* (PB) [Act-RFP], a PB-transposon, into the FVB/N background (48). Hybrid (C57BL/6 X CBA/He) mice in some experiments were housed and bred in the Gore Hill Research Laboratory, St Leonards, New South Wales, Australia. Experiments were performed at the Shanghai Institute of Planned Parenthood Research, Wenzhou Medical University and the Kolling Institute, University of Sydney.

### Embryo collection and culture

Six-week-old females were superovulated by intraperitoneal injection of 5 IU pregnant mare serum gonadotropin (PMSG) (Ningbo Second Hormone Factory, China). After 48 h, mice were injected again with 5 IU human chorionic gonadotrophin (hCG) (Livzon, Zhuhai, China). Pregnancy was confirmed by the presence of a copulation plug the following morning. Oocytes or zygotes were recovered 20 h post-hCG from unmated and mated females, respectively. Cumulus cells were removed by brief exposure to 300 IU hyaluronidase (Sigma Chemical Company, St Louis, MO, USA). Two-cell, 4-cell, 8-cell embryos, and blastocysts were isolated from the oviducts and/or uterus of plug-positive female mice at 40, 60, 68 and 90 h after hCG injection, respectively. Oocytes and embryos were collected in HEPES-buffered modified human tubal fluid medium (HEPES-HTF) (49). Embryos were cultured at density of 1 or 10 embryos in 10 μl KSOM medium (50) according to the experimental design in 60-well plates (LUX 5260, Nunc, Naperville, IL, USA) overlaid with 2 mm of heavy paraffin oil (Sigma) at 37°C in 5% CO2 under 5% O2 or 20% O2 tension. All components of the media were tissue culture grade (Sigma) and contained 3 mg bovine serum albumin/mL (Sigma). Treatments were performed in KSOM medium supplemented with: (1) 1, 10, 100, or 1000 nM of 2-(4-amino-1-isopropyl-1H-pyrazolo[3,4-d]pyrimidin-3-yl)-1H-indol-5-ol (PP242; Sigma, P0037), (2) 2, 10, or 50 μM 4EGI-1 (Merck, Germany), (3) 10 μM SB203580 (16, 28) (Merck), (4) 25 nM staurosporine (28) or amino acids (51) (Sigma).

### Sperm samples

Mouse sperm was collected from the epididymis of 10 to 12-week-old hybrid males. Sperm were prepared in in pre-equilibrated HTF (52) and then centrifuged for 10 min, 500×g at room temperature (RT). The supernatant was carefully decanted and the sperm pellet was resuspended in PBS for immunofluorescent procedures.

### Monitoring of RFP reporter expression

*In vitro* embryos expressing the RFP reporter were monitored with an inverted epi-fluorescent microscope (Nikon, Japan) at 24 h intervals. Baseline fluorescence was set as the measured fluorescence in wild type embryos.

### Genotyping

Genomic DNA was extracted from mouse tails and individual embryos with 180 μL or 5 μL50 mM NaOH at 95°C 10 min, and 20 μL or 0.5 μL 1M Tris-HCl (pH 8.0), respectively. DNA was used as templates for PCR (KOD-FX, TOYOBO, Osaka, Japan) with the primer pairs: *PB* primer (RF1) CCTCGATATACAGACCGATAAAACACATGC, GL-primer TGCTTATCAACAAAAAGCAGATGGC, and GR-primer ACAGGAAAGGAGACAGTACCTGAG. Insertion band size was 580 bp and GL/GR 949 bp. PCR amplification products were analyzed by electrophoresis on 2% (w/v) agarose gel staining with SYBR green to visualize PCR products on u.v. transilluminator. Fragments were verified by size and representative samples had sequence analyzed (Shanghai Jieli Science and technology Ltd.co, China).

### IVF and ICSI

Concentrated sperm was collected from the epididymis of 10 to 12-week-old hybrid males and incubated in pre-equilibrated HTF(52) for 15-30 min at 37°C with 5% CO2. Cumulus-oocyte complexes (COC) were collected from oviducts of female hybrids 13-15 h post-hCG and briefly treated with hyaluronidase HEPES-HTF medium. COC were washed and moved into HTF medium in preparation for intracytoplasmic sperm injection (ICSI).

Sperm was observed under an Olympus IX75 microscope (Olympus, Hamburg, Germany). Fresh sperm were incubated in drops of equilibrated HTF for 1 h at a final concentration of 1 × 10^6^/mL. COC were added and incubated for 6 h. Fertilized oocytes at pronuclear stage were picked out and transferred to 20 μL KSOM medium drops (20 embryos per drop).

For ICSI procedure, 1 μL of fresh sperm was mixed with a drop of HEPES-HTF medium containing 10% (w/v) polyvinylpyrrolidone (PVP, Sigma). Piezo pulses were used to separate the sperm head from the tail, so the head could be injected into the oocyte. The embryos were observed for pronuclear embryos, 2-cell embryos, and blastocysts at 6 h, 24 h, and 96 h after insemination, respectively.

### Blastocyst outgrowth

Blastocysts developed *in vitro* were cultured in each well of a 24-well plate with 1 mL Dulbecco’s modified Eagles medium (DMEM, Sigma) supplemented with 10% (v/v) heat inactivated (40 min at 56°C) fetal bovine serum (FBS, Sigma), and antibiotics (0.6 mg penicillin/CSL/mL, 1 mg streptomycin/mL; Sigma). After 96 h, the embryos were analyzed with genotyping and immunofluorescence.

### Immunofluorescence

Immunofluorescence was performed as previously described (28, 53). After fixation, permeabilization and blocking, embryos or sperm were incubated overnight at 4°C with primary antibodies: 2 μg/mL rabbit anti-eIF4E polyclonal IgG, 2 μg/mL rabbit anti-p-eIF4E (S209) polyclonal IgG, 2 μg/mL mouse anti-4E-BP1 polyclonal IgG, 2 μg/mL rabbit anti-p4E-BP1+2+3 (T45) polyclonal IgG, 2 μg/mL rabbit anti-OCT3/4 polyclonal IgG, 2 μg/mL rabbit anti-p-S6K1 (T389) polyclonal IgG, anti mouse mTOR monoclonal IgG, anti-rabbit p-mTOR(ser2448) polyclonal IgG, and 2 μg/mL isotype negative control immunoglobulin. All primary antibodies were purchased from Abcam (Cambridge, MA, USA). Primary antibodies were detected by Texas Red-conjugated goat-anti mouse (Sigma) or FITC-conjugated goat anti-rabbit (Sigma) secondary antibodies for 1 h at room temperature. Whole section imaging was performed by mercury lamp UV illumination with an epifluorescent Nikon ECLIPSE 80i microscope, using a Plan Apo 40X/1.0 oil objective. Optical sectioning was performed with a Nikon A1+ confocal microscope (Nikon, Tokyo, Japan) equipped with a Plan Apo 60X oil objective. All quantitative analysis of immunofluorescence experiments was performed with Image-Pro Plus (version 6.3, Media Cybernetics USA).

### Western blot

Western blot analysis was performed as previously described (53). The embryos were washed in cold PBS, lysed in extraction buffer, followed by three cycles of freezing and thawing in liquid nitrogen and vortexing, respectively. The extracted embryo proteins were diluted with Laemmli loading buffer, separated on 20% homogenous SDS-polyacrylamide gels (GE Healthcare Australia Pty. Ltd. Rydalmere, New South Wales, Australia) using a PhastSystem workstation (Amersham Pharmacia Biotech, Sweden). Proteins were then transferred onto polyvinylidene fluoride membrane (PVDF, Hybond-P, Amersham) in transfer buffer containing 12 mM Tris (Sigma), 96 mM glycine (BDH, Sydney, Australia), and 20% (v/v) methanol (BDH) by a semi-dry PhastTransfer system (Amersham). The membrane was incubated overnight at 4°C in 10 mL of blocking buffer containing 2.5% (w/v) skim milk powder (Diploma, New Zealand) with 0.2 μg/mL primary antibodies. The membrane was then incubated in 1:5000 horse radish peroxidase (HRP) conjugated secondary antibody (Sigma), followed by Super Signal^@^ West Femto chemiluminescent substrate (Pierce, Rockford, IL, USA) and the products were recovered by Laser 4000 chemiluminescent system (Bio-Rad, CA, USA). Finally, the membrane was stripped by incubation in 200 mM NaOH (Sigma) for 30 min at room temperature and re-probed with 1:2000 rabbit anti**-** ACTIN IgG (Sigma).

### Statistical analysis

Statistical analysis was performed with SPSS for Windows (Version 22.0, SPSS Inc., Chicago, IL, USA). Fluorescence intensity, optical intensity, area, and cell number were quantitatively analyzed by univariate analysis of variance. Those parameters were set as the dependent variable, while the test treatments were the independent variables. Experimental replicates were incorporated into the model as covariates. Differences between individual treatments were analyzed by the least significance difference test. Blastocyst development rate was assessed by binary logistic regression analysis. Chi-square tests were used to determine the differences between observed and expected frequency distributions. Less than a 5% probability (P < 0.05) was considered significant.

## Supporting information

Li-Manuscript

## Acknowledgments

We thank Nanjing Your Bio-tech Development Ltd. Co (Jiangbei New District, Nanjing, Jiangsu Province, China) for generous donating all embryonic culture media; KSOM, HEPES-HTF and HTF.

This work was supported by grants from the National Natural Science Foundation of China awarded to X.J (81471458), and the Australian National Health and Medical Research Council (NHMRC) awarded to C.O.

