## Supplementary material for "The regulation of mammalian maternal-to-embryonic transition by Eukaryotic translation initiation factor 4E": Li-Manuscript

Dr. Xingliang Jin,

**This PDF file includes:**

Figures S1
Tables S1

Suppl Fig 1

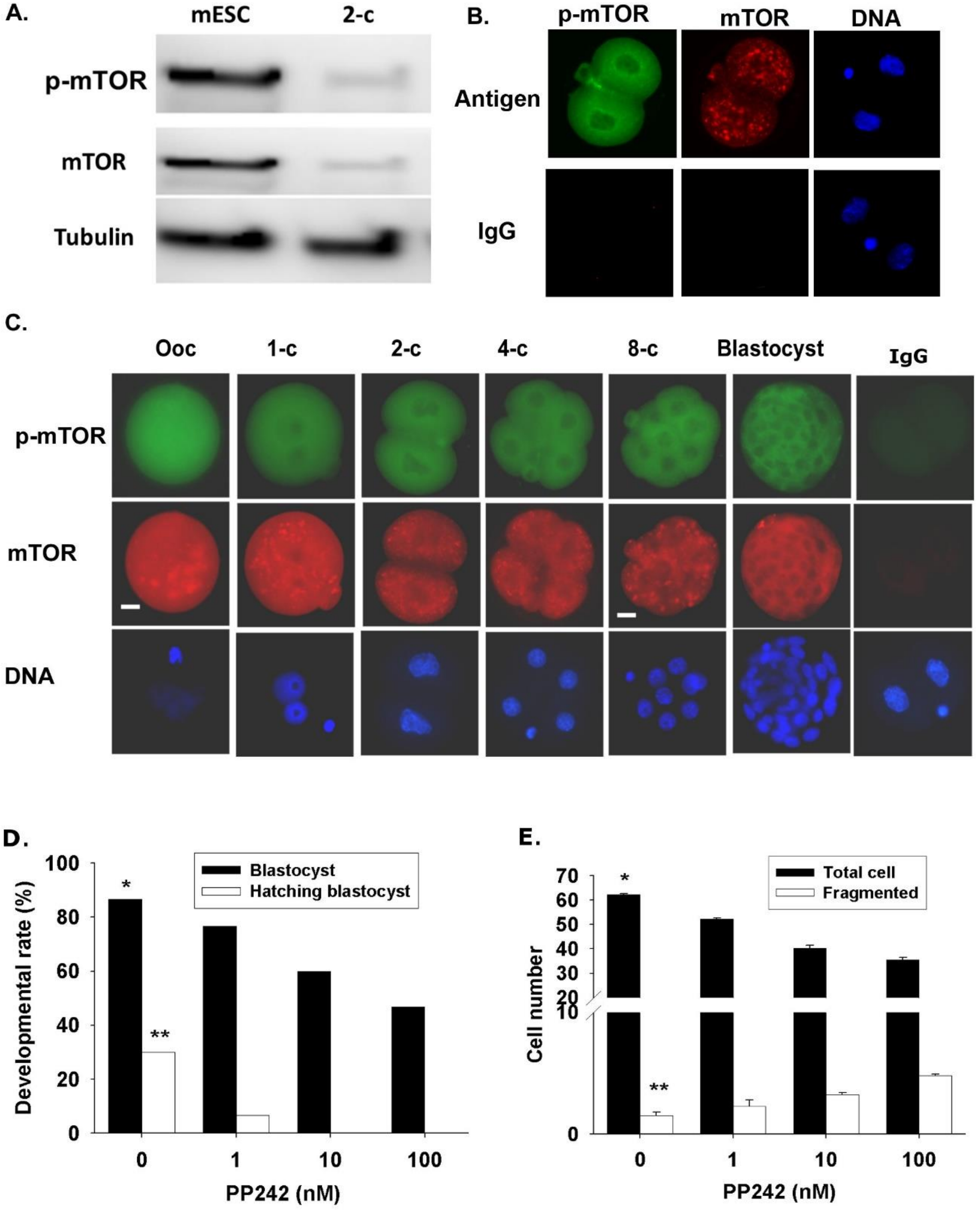


**Fig. S1 mTOR and p-mTOR in mouse preimplantation embryos.**

mTOR and p-mTOR were analyzed in mouse 2-cell embryo by western blot (A). Triple Immunostaining of mTOR, p-mTOR, and Hoechst 33342 DNA in a 2-cell embryo imaged by confocal microscope (B) and in preimplantation development by epi-fluorescent microscope (C). The data were representative of three independent replicates with 10 embryos analyzed for each developmental stage. Scale bar = 10 mm for all images. (D) The data were representative of three independent replicates performed to analyze the effects of PP242 on the rate of blastocyst formation and hatching blastocysts, as well as, the number of total and fragmented cells in formed blastocysts. Each treatment included at least 20 embryos in each replicate. *P < 0.001 and **P < 0.01, comparing to PP242-treated groups.

### Table S1.

**Table S1 the number of embryos and RFP+ embryos in developmental landmarks**

| | No. of Mating female | No. of virginal plug | Zygotes at 20 h post-hCG | | | 2-cell embryo at 40 h post-hCG | | | 4-8 cell embryo at 64h post-hCG | | |
|---|---|---|---|---|---|---|---|---|---|---|---|
| | | | Mean ± S.E.M | total | | Mean ± S.E.M | total | | Mean ± S.E.M | total | |
| | | | | No | RFP+ | | No | RFP+ | | No | RFP+ |
| ***Eif4e*$^{+/-}$ ♂ × *Eif4e*$^{+/-}$ ♀** | 90 | 37 (31.1%)* | 15.4 ± 1.1 | 202 | 96 (47.5%) | 12.7 ± 0.8** | 151 | 0 (0%) | 11.9 ± 1.4* | 64 | 44 (68.8%) |
| ***Eif4e*$^{+/+}$ ♂ × *Eif4e*$^{+/-}$ ♀** | 25 | 23 (92.0%) | 16.3 ± 0.8 | 147 | 72 (49%) | 15.8 ± 1.1 | 110 | 58 (52.3%) | 16.3 ± 0.9 | 134 | 65 (48.5%) |
| ***Eif4e*$^{+/-}$ ♂ × *Eif4e*$^{+/+}$ ♀** | 29 | 25 (86.2%) | 16.2 ± 1.1 | 80 | 0 (0%) | 16.1 ± 0.9 | 81 | 0 (0%) | 16.1 ± 1.3 | 90 | 72 (40.0%) |
| ***Eif4e*$^{+/+}$ ♂ × *Eif4e*$^{+/+}$ ♀** | 20 | 18 (90%) | 20.2 ± 0.8 | 165 | NA | 18.3 ± 0.6 | 183 | NA | NA | | |

Discovering plugs in female virginals indicate fertilization. Its rate, the number of embryos and RFP$^{+}$ embryos were shown in several developmental stages that freshly collected from female reproductive tracts. *p<0.001 and **p <0.01, compared to the according to *Eif4e*$^{+/+}$ ♂ × *Eif4e*$^{+/+}$ ♀ mating or *Eif4e*$^{+/-}$ ♂ × *Eif4e*$^{+/+}$ ♀ mating *and Eif4e*$^{+/+}$ ♂ × *Eif4e*$^{+/-}$ ♀.
